# Sub-phenotypes of pneumonia defined by pulmonary histopathological features

**DOI:** 10.1101/2025.05.18.654716

**Authors:** Amulya Shastry, Bradley E. Hiller, Nathan L. Sanders, Anna E. Tseng, Jeet Kothari, Aoife K. O’Connell, Hans P. Gertje, Catherine T. Ha, Ekaterina Murzin, Katrina E. Traber, James A. Lederer, Mattthew R. Jones, Florian Douam, Thomas G. Beach, Daniel G. Remick, Nicholas A. Crossland, Stefano Monti, Joshua D. Campbell, Joseph P. Mizgerd

## Abstract

Establishing sub-phenotypes of pneumonia based on distinct host processes will be a step towards using host-directed therapies (to complement microbe-directed therapies) more rationally and precisely. Although pneumonia is a pulmonary pathophysiology, histological changes within the lungs have not been leveraged for sub-phenotyping. We addressed this by scoring 18 histopathology features (e.g., type 2 cell hyperplasia or necrosis) across rapid autopsy lung samples from 276 elderly subjects with pneumonia. Machine learning algorithms segregated subjects into seven different sub-phenotypes of pneumonia with distinct histopathology signatures. Quantitative immunofluorescence demonstrated associations of macrophages, neutrophils, T cells, and B cells with select histology features and pulmonary pathology sub-phenotypes. Mouse models revealed corollary sub-phenotypes, although some histology features observed in human lungs were never observed in mice. By illuminating this spectrum of histopathologies and discriminating discrete sub-phenotypes of pneumonia, a foundational framework emerges for developing and using host-directed therapies for subsets of pneumonia patients.

## Introduction

Pneumonia causes a tremendous burden of disease worldwide, particularly due to childhood deaths in lower income countries^1^. In wealthier countries like the United States, pneumonia is the leading cause of death due to infection^2^ because of higher mortality rates among older adults, with a higher risk of death than any other common cause of hospitalization for those over 65 years of age^3,4^. Beyond deaths, pneumonia causes substantial morbidity including post-acute sequelae that accelerate disease processes and unhealthy aging^3,5^.

Pneumonia results from lower respiratory infections that stimulate diverse biological processes yielding pulmonary infiltrates. A wide variety of microbes cause pneumonia, including bacteria (e.g., *Streptococcus pneumoniae*, *Legionella pneumophila, Klebsiella pneumoniae*), viruses (e.g., influenza viruses, coronaviruses, respiratory syncytial virus), and fungi (e.g., *Cryptococcus neoformans*, *Pneumocystis jirovecii*, *Aspergillus fumigatus*)^3,6–9^. Microbiological and molecular tests are improving^10–12^, but pneumonia etiology usually remains uncertain^7,13–15^. Etiological inferences can be refined by infection characteristics, such as where it was contracted (e.g., community-acquired or hospital-acquired) or is situated (e.g., lobar pneumonia or empyema). Assessments of infectious etiology guide the use of microbe-directed therapies like antibiotics, antivirals, and antifungals.

In contrast, host-directed therapies are not directed to relevant patients based on the nature of host responses^3,6,16,17^. Many host-directed therapies have been proposed for pneumonia patients (corticosteroids, anti-inflammatories, biologics inhibiting cytokine activities, recombinant immunostimulatory proteins, coagulation modulators, statins, antioxidants, etc.)^16,18^. Corticosteroids have been most extensively investigated, with demonstrated utility for some settings of pneumonia^19^, but which patients may respond remains uncertain. Defining sub-phenotypes of pneumonia based on host responses will facilitate precision medicine by improving the development and rationally guided delivery of host-directed therapies^16,17,20^.

While biological sub-phenotypes have not yet been established for pneumonia, there has been progress in this area for two important acute sequelae of pneumonia, ARDS^21,22^ and sepsis^23^. Patients with ARDS or sepsis can be sub-phenotyped into hyperinflammatory and hypoinflammatory sub-phenotypes based primarily on blood analytes, discriminating individuals likely to respond differently to host-directed therapies^24^. However, blood sample measurements are often discordant with ongoing pulmonary processes^25^. Examination of pulmonary pathology for ARDS patients reveals that only about half demonstrate the hallmark feature of diffuse alveolar damage (DAD)^26–28^. Compared to ARDS patients without DAD, those with DAD have worse pulmonary physiology^28^ and an increased risk of death^27^, with death more likely from hypoxemia and less likely from shock^28^. A small fraction of pneumonia patients develops ARDS, which includes distinct pulmonary histopathological presentations (DAD or not) that have physiological and prognostic significance. While pneumonia is a pulmonary pathophysiology, pneumonia patients have not been sub-phenotyped based on biological processes within the lungs. Focusing on the lungs, where the pneumonia process is occurring, we endeavored to determine whether distinct sub-phenotypes of pneumonia would emerge based on variations in local inflammation and damage revealed by pulmonary histopathology.

## Results

### Human rapid autopsy samples as a pneumonia-relevant biobank for analyzing lung histopathology

To compile a pneumonia-relevant histology biobank, retrospective tissues were collected from subjects in the greater Phoenix, AZ, retirement communities who consented for research-purpose rapid autopsies between the years of 2006 and 2020^29^ (**Fig. 1A**). We included 361 subjects who received a diagnosis of acute pneumonia upon autopsy prior to 2020, 25 from 2020 with COVID-19 based on SARS-CoV-2 RT-PCR nasopharyngeal swab, and 18 from subjects from those years with no known lung disease. All samples were stained by H&E and anti-fibrinogen immunohistochemistry (IHC). After excluding samples with pulmonary infarction or concurrent neoplastic process, this retrospective cohort provided 292 cases for analysis: 253 from patients with a pneumonia diagnosis prior to 2020 (hence not COVID), 23 with pneumonia and a COVID-19 diagnosis, and 16 “control” samples from patients with no visible lung disease upon autopsy and no known history of lung disease (**Fig. 1B**). Before inspecting the samples, we selected 20 histopathological features relevant to pneumonia for examination (**Supp. Fig. 1A**, with examples shown in **Fig. 1C-K** and **Supp. Fig. 1B-L**). For each feature, two board-certified pathologists provided a score of 0 (not observed), 1 (present in <5% of the section), 2 (present in 5-25% of the section), or 3 (present in >25% of the section). Two features (vasculitis and hemophagocytosis) were observed in fewer than 10 samples and removed from further analyses.

**Figure 1.**
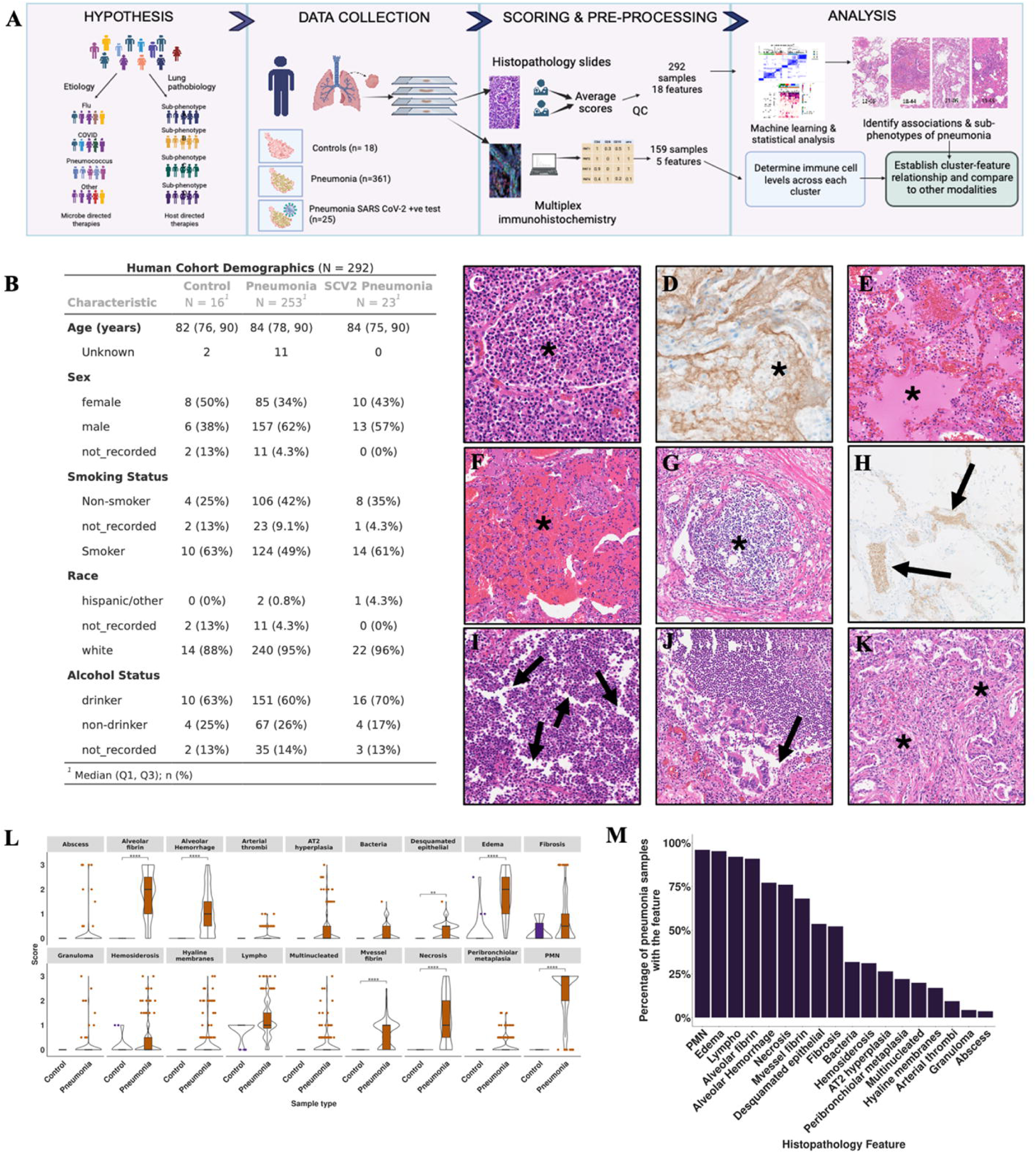
Histopathological features of pneumonia, from human rapid autopsy samples. (A-B) To identify lung pathobiology sub-phenotypes, we utilized rapid autopsy lung sections from subjects diagnosed with pneumonia at autopsy. These samples were stained by H&E, anti-fibrinogen IHC, and qmf-IHC, independently scored by two pathologists, and analyzed computationally. (C-K) Representative images of the 9 most frequent pneumonia histopathology features observed in human samples, including neutrophils (C, H&E), alveolar fibrin (D, anti-fibrinogen IHC), edema (E, H&E), alveolar hemorrhage (F, H&E), lymphoplasmacytosis (G, H&E), microvessel fibrin (H, anti-fibrinogen IHC), necrosis (I, H&E), desquamated epithelia (J, H&E), and fibrosis (K, H&E). (L) Scores for each histopathology feature were compared between the control samples with no known lung disease and the pneumonia samples. Data were represented as median ± SEM and analyzed by Kruskal-Wallis test, followed by a post-hoc Dunn test and adjusted for multiple tests using Bonferroni correction method. Asterisks show adjusted p-values with the following range: 0-1e-04(****), 1e-04 - 0.001(***), 0.001-0.01(**), 0.01-0.05(*), 0.05-1 (ns). (M) Frequencies of features binarized as present or absent across the cohort of pneumonia samples.

### Associations of histopathological features with pneumonia and its etiology

The control (non-pneumonic) lung samples had incidental findings consistent with the advanced age of this cohort, including low levels of edema, fibrosis, hemosiderosis, and lymphoplasmacytosis (**Fig. 1L**). The pneumonia samples had significantly higher mean scores for multiple features compared to the control group, including neutrophils (2.3 ± 0.9), edema (1.8 ± 0.8), alveolar fibrin (1.8 ± 0.9), necrosis (1.3 ± 1.0), alveolar hemorrhage (1.1 ± 0.9), microvessel fibrin (0.7 ± 0.6), and desquamated epithelia (0.3 ± 0.3) (**Fig. 1L**). This collection of features may therefore be considered characteristic of pneumonia. However, remarkable variation was noted in all features, and no single feature was consistently observed in every pneumonic sample (**Fig. 1M**).

Histopathology features may not present independently, for example if they involve shared or inter-dependent processes. To determine associations in pneumonic lungs, we assessed all feature-by-feature relationships using a Spearman correlation metric. The strongest positive relationship was between neutrophils and necrosis (**Fig. 2A**). Neutrophils and necrosis also consistently correlated with edema, alveolar hemorrhage, alveolar fibrin, microvessel fibrin, and desquamated epithelia (**Fig. 2A**), A separate set of features showed consistent associations with each other, including lymphoplasmacytosis, fibrosis, alveolar epithelial type 2 cell (AT2) hyperplasia, peribronchiolar metaplasia, and multinucleated cells (**Fig. 2A**). Lymphoplasmacytosis, fibrosis, and AT2 hyperplasia all showed consistent negative correlations with necrosis, neutrophils, edema, and alveolar hemorrhage. These systematic analyses suggest that pneumonic lungs include at least 2 discrete patterns of histopathological presentations.

**Figure 2.**
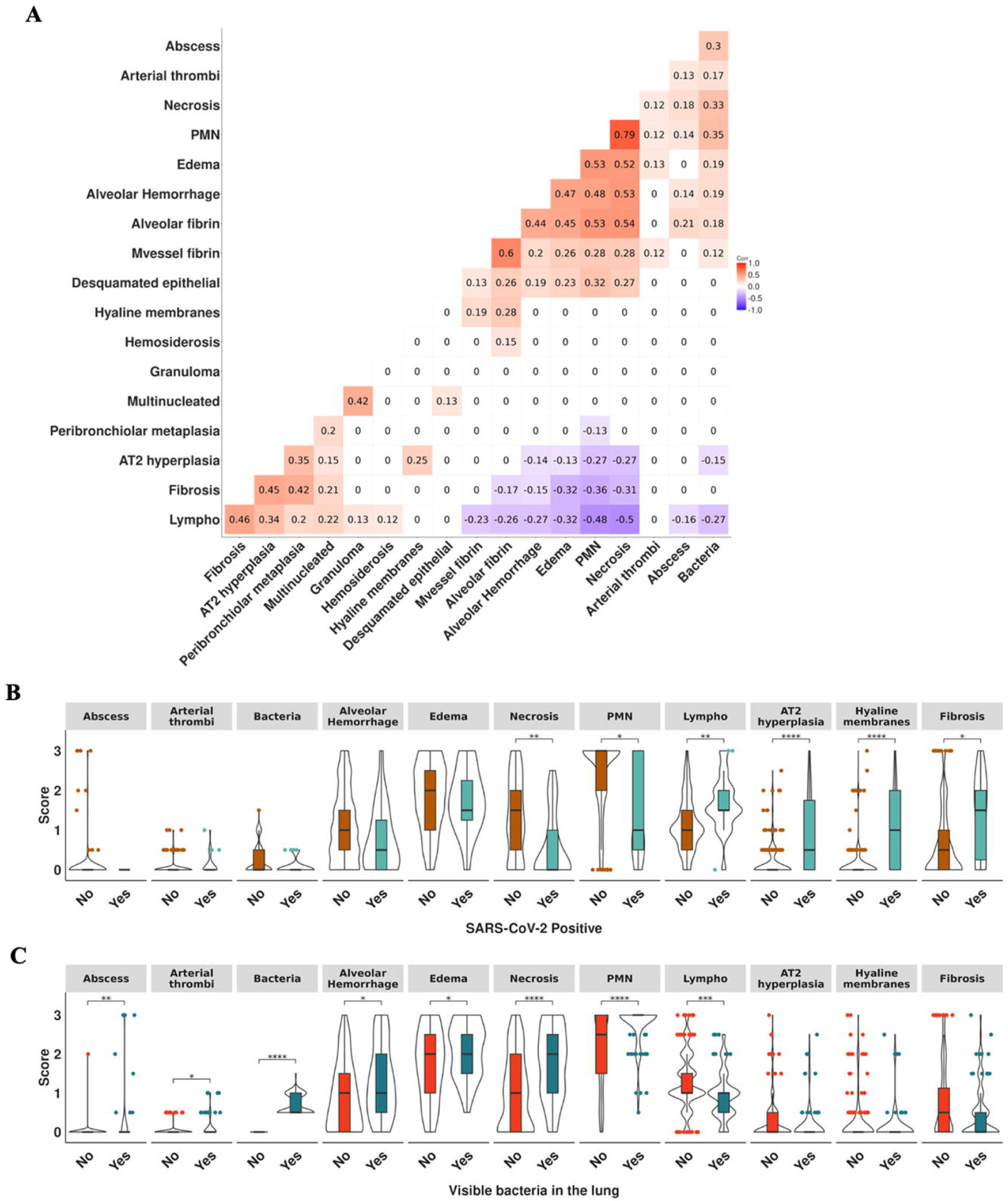
Associations amongst histopathological features in pneumonic human lungs. (A) A Spearman correlation plot across histopathological features. Correlations reaching statistical significance were communicated in the figure as correlation coefficients ranging from - 1.0 to 1.0, while non-significant correlations were assigned a 0 value. To test significance, two tailed Wilcoxon-test was used with BH correction for multiple tests. (B-C) Histopathology feature scores were compared in pneumonia samples with (B) a SARS-CoV-2 diagnosis or (C) with visible bacteria in the H&E section (B) against all other pneumonia samples. Data were represented as median ± SEM and analyzed by Kruskal-Wallis test, followed by a post-hoc Dunn test and adjusted for multiple tests using Bonferroni correction method.

Etiology is usually uncertain for pneumonia^7,15^, which also applied to subjects in our dataset. Two types of data were leveraged to infer microbial etiology for some samples: SARS-CoV-2 diagnosis by nasal sample RT-PCR (suggesting these pneumonias may be due to that virus) and the presence of visible bacteria in the H&E-stained section (suggesting those pneumonias may be due to bacterial infection). When compared to all other pneumonia samples, the SARS-CoV-2 samples exhibited significantly higher AT2 hyperplasia and hyaline membranes (**Fig. 2B**), which together form typical features of DAD and have been associated with SARS-CoV-2 pneumonia^30–32^. Additionally, SARS-CoV-2 samples had significantly higher fibrosis and lymphoplasmacytosis and significantly less necrosis and neutrophils compared to the other pneumonias (**Fig. 2B**). Other features were not significantly different (**Fig. 2B, Supp. Fig. 2A**). To investigate whether the observation of bacteria correlated with pulmonary histopathology, we binarized the bacterial feature to Yes/No. Sections with bacteria had significantly higher scores for neutrophils, necrosis, edema, abscess, arterial thrombi, and alveolar hemorrhage compared to samples without visible bacteria (**Fig. 2C**). In addition, they had less lymphoplasmacytosis than samples without bacteria (**Fig. 2C**). There were no significant differences in other features (**Fig. 2C, Supp. Fig. 2B**). These analyses suggest different etiologies associate with different pulmonary histopathologies. A collection of features including AT2 hyperplasia, hyaline membranes, lymphoplasmacytosis, and fibrosis associate with each other across pneumonic lungs, which may be characteristic of COVID pneumonia. A different collection of features including necrosis, neutrophils, edema, alveolar fibrin, and alveolar hemorrhage associate with each other across pneumonic lungs, which may be characteristic of bacterial pneumonias.

### Machine learning distinguishes seven pulmonary pathology sub-phenotypes of pneumonia

To better understand heterogeneity in pneumonia, we employed unsupervised machine learning using the 18 histopathology features to cluster pneumonia subjects into sub-phenotypes (**Fig. 1A**). After using consensus clustering with distances-based and variation-based metrics to determine the optimal number of clusters^33^ (**Supp. Fig. 3A-C**), we identified seven clusters of pneumonia samples distinguished by specific histopathological signatures (**Figure 3A-C, Supp. Fig. 3A-C**).

**Figure 3.**
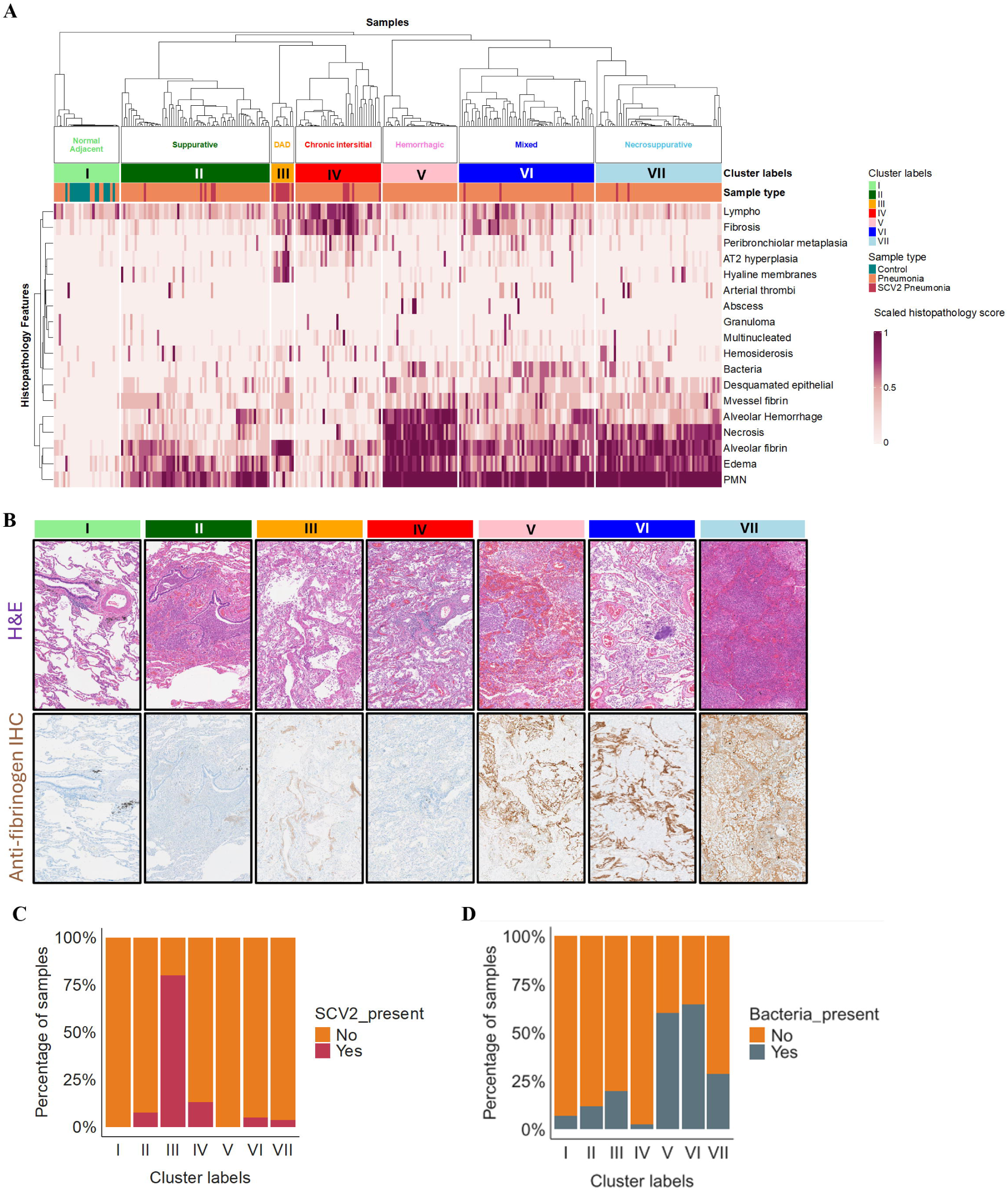
Sub-phenotypes of pneumonia based on pulmonary histopathology revealed by human rapid autopsy samples. (A) Based on 18 scored features (rows) for each of 292 human subjects (columns), clustering metrics identified 7 subsets of pneumonia (I-VII) that were distinguished by pulmonary histopathology. (B) Representative images of H&E-stained and anti-fibrinogen IHC-stained sections for each of the pneumonia histopathology sub-phenotypes. (C-D) Samples were binarized for (C) SARS-CoV2 positive PCR test (in C) or presence of visible bacteria in the lung (in D).

All 16 “no pneumonia” controls (100%) as well as 13 of the pneumonia samples (4.7%) were grouped in Cluster I (**Figure 3A-B**). These samples displayed moderate levels of lymphoplasmacytosis, with typically low but variable and inconsistent levels of other features. We named this cluster “normal adjacent.” Cluster II (24% of c elevated scores for neutrophils and edema (**Fig. 3A-B**), which we named “Suppurative” pneumonia.

Cluster III (3.6%) was characterized by coincident hyaline membranes and AT2 hyperplasia and thus named “DAD.” (**Fig. 3A-B**). In addition, this cluster had high alveolar fibrin despite little or no necrosis, as well as consistent fibrosis and lymphoplasmacytosis. Notably, 8 of these 10 samples were SARS-CoV-2+ cases (**Fig. 3C**), consistent with AT2 hyperplasia and hyaline membranes being hallmarks of severe COVID in 2020^30–32^. Cluster IV (12%) also had elevated lymphoplasmacytosis and fibrosis, but it lacked the higher levels of hyaline membranes, AT2 hyperplasia, or alveolar fibrin observed in Cluster III (**Fig. 3A-B**). In addition, Cluster IV had lower neutrophils, alveolar fibrin, necrosis, and edema than most other clusters. We named this cluster “chronic interstitial.”

Clusters V, VI, and VII all had consistently high neutrophils and alveolar fibrin (**Fig. 3A-B**). Together, these three clusters composed 54% of the pneumonia samples. Cluster V (12%) was characterized by consistently high alveolar hemorrhage and edema in addition to neutrophils, alveolar fibrin, and necrosis, so we named it “hemorrhagic.” Cluster VI (22%) was the most internally heterogeneous cluster, with consistently high neutrophils and alveolar fibrin but edema that was less prominent than in clusters V and VII and other features more varying (**Fig. 3A**). One subset within this cluster, segmented to the left, had increased fibrosis and lymphoplasmacytosis in addition to abundant neutrophils and alveolar fibrin (**Fig. 3A**). Another more central subset had particularly high bacteria (**Fig. 3D**) along with neutrophils and alveolar fibrin, but little necrosis. The third subset, segmented to the right, had high alveolar hemorrhage without the concomitant necrosis and edema characteristic of other samples with prominent hemorrhage. We named Cluster VI “Mixed” to reflect this heterogeneity. Cluster VII (20%) showed prominent necrosis in the lung parenchyma in addition to high neutrophils, edema, and alveolar fibrin (**Fig. 3A-B**). We named this “Necrosuppurative.”

Variable importance scoring revealed that each pneumonia histopathology cluster was associated with distinct sets of driving features that were especially low or high in that cluster compared to others (**Supp. Fig. 3D**). Age, sex, smoking status, and alcohol consumption did not associate selectively with any of the different pneumonia features or clusters (**Supp. Fig. 4-5**). We propose that these clusters represent sub-phenotypes of pneumonia, differentiated by distinct pulmonary histopathology patterns.

### Associations of leukocytes with pneumonia histopathologies and sub-phenotypes

To determine whether immune cells associated with observed histopathological features or with the sub-phenotypes derived from clustering, we used quantitative multiplex fluorescence immunohistochemistry (qmf-IHC) to determine the relative content of myeloperoxidase-positive (MPO^+^) cells (interpreted as neutrophils), CD68^+^ cells (interpreted as macrophages), CD19^+^ cells (interpreted as B cells), CD4^+^ cells (interpreted as helper T cells), and CD8^+^ cells (interpreted as cytotoxic T cells) (**Fig. 4A**). Compared to control samples, the pneumonia samples had significantly higher macrophages and neutrophils, but none of the lymphocytes reached significance (**Supp Fig. 6A**). Compared to non-SARS-CoV-2+ pneumonias, SARS-CoV-2+ samples had significantly increased cytotoxic T cells and reduced neutrophils (**Supp Fig. 6B**). There were no significant differences in the abundance of these leukocytes between those pneumonia samples with *vs.* without observed bacteria (**Supp Fig. 6C**).

**Figure 4.**
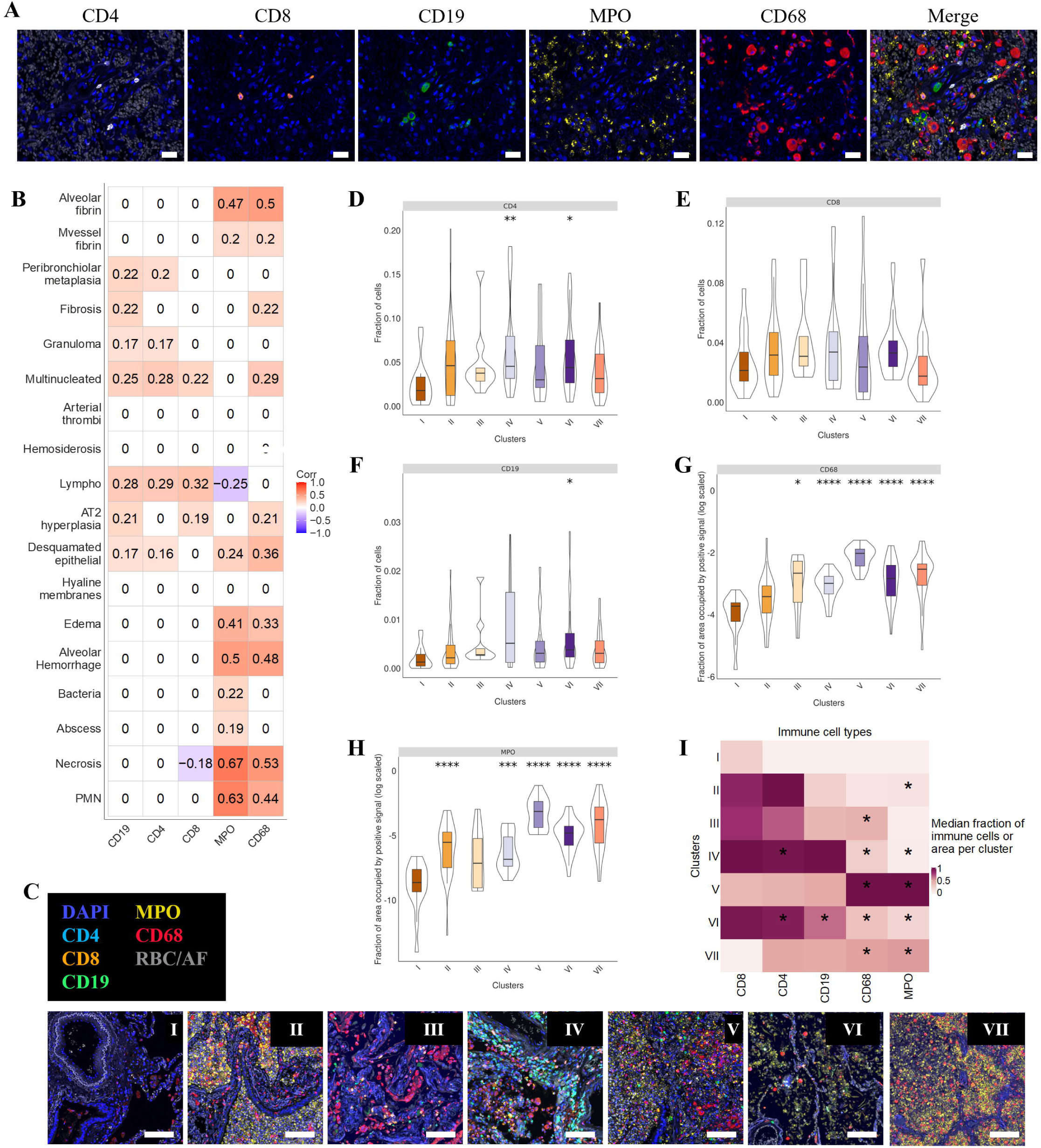
Immune cell associations with histopathology features and sub-phenotypes in pneumonic human lungs. (A) qmf-IHC was used to identify CD4 (white), CD8 (orange), CD19 (green), MPO (yellow), and CD68 (red) in each human rapid autopsy sample of sufficient quality for staining. Scale bars represent 100 µm. (B) A Spearman correlation plot defined the associations between each pneumonia histopathology feature and qmf-IHC target. Significant correlations were reported as correlation coefficients from -1.0 to 1.0 and non-significant correlations were assigned a 0 value. (C) Representative images of qmf-IHC staining in each pneumonia sub-phenotype cluster. Scale bars represent 100 µm. (D-H) Using cluster I as a hanging control, positive intensity signals were compared to all other clusters across CD4, CD8, CD19, CD68, and MPO positive cells. To determine significance, the two tailed Wilcoxon-test was used with Bonferroni correction for multiple tests. Asterisks show adjusted p-values with the following range: 0-1e-04 (****), 1e-04 - 0.001(***), 0.001-0.01(**), 0.01-0.05(*), 0.05-1 (ns). (I) Heatmap of the median intensity per cluster group across the 5 immune cell signals, with asterisk indicating significance at P<0.05 as calculated in D-H.

To determine how immune cells associate with distinct histopathology features, we calculated Spearman correlation coefficients for each pairing (**Fig. 4B**). As anticipated, B cells, helper T cells, and cytotoxic T cells each significantly positively correlated with lymphoplasmacytosis, as did neutrophils scored by pathologist inspection with neutrophils quantified by MPO qmf-IHC (**Fig. 4B**). In addition, B cells positively associated with multinucleated cells, peribronchiolar metaplasia, fibrosis, AT2 hyperplasia, granuloma, and desquamated epithelia (**Fig. 4B**). Helper T cells positively associated with multinucleated cells, peribronchiolar metaplasia, granuloma, and desquamated epithelia (**Fig. 4B**). Cytotoxic T cells correlated positively with multinucleated cells and AT2 hyperplasia, but negatively with necrosis (**Fig. 4B**). Macrophage and neutrophil staining both associated with necrosis, alveolar and microvessel fibrin, alveolar hemorrhage, edema, and desquamated epithelia (**Fig. 4B**). In addition, neutrophils associated positively with bacteria and abscess, and negatively with lymphoplasmacytosis, while macrophages associated with multinucleated cells, fibrosis, and AT2 hyperplasia (**Fig. 4B**). Some features (arterial thrombi, hemosiderosis, and hyaline membranes) did not significantly associate with any of the 5 leukocytes examined.

Using these qmf-IHC data, we determined whether and which immune cells were significantly elevated in the pneumonia histopathology sub-phenotypes compared to cluster I (Normal Adjacent). Each immune cell type exhibited unique pneumonia sub-phenotype combinations (**Fig. 4C**). Helper T cells were significantly increased in clusters IV (chronic interstitial) and VI (Mixed) (**Fig. 4D**). Cytotoxic T cells did not differ (**Fig. 4E**). B cells were significantly increased in cluster VI (Mixed) (**Fig. 4F**). Macrophages were significantly elevated in clusters III (DAD), IV (chronic interstitial), V (hemorrhagic), VI (Mixed), and VII (necrosuppurative) (**Fig. 4G**). Neutrophils were significantly increased in clusters II (suppurative), IV (chronic interstitial), V (hemorrhagic), VI (Mixed), and VII (necrosuppurative) (**Fig. 4H**). To compare immune cell patterns across clusters, we calculated median fraction of immune cells (T and B cells) or median fraction of area (macrophages and neutrophils) per cluster and min-max scaled the values per cell type. Each sub-phenotype of pulmonary pathology displayed a distinct immune cell signature (**Fig. 4I**).

### Pneumonia sub-phenotypes in mouse models

While mouse models are frequently used to study pneumonia, no studies have systematically compared histopathological features from infected mouse lungs to pneumonic human lungs^34,35^. We aimed to determine which features and sub-phenotypes of human pneumonia are reflected or missed in commonly used mouse models of pneumonia. We characterized pulmonary histopathology in 100 murine lung samples derived from 9 mouse models of pneumonia (and 2 sets of controls) in the same manner as with our human lung samples (**Fig. 5A**). Young adult mice (8-14 weeks old) were challenged with saline, bacterial pathogens (*Streptococcus pneumoniae* serotype 3 [*Sp3*] or serotype 19F [*Sp19F*], *Escherichia coli*, or *Klebsiella pneumoniae*), or viral pathogens (the PR8 strain of H1N1 influenza A virus or mouse-adapted strains of SARS-CoV-2, MA10 or MA30) at doses sufficient to be lethal in both male and female mice (**Fig. 5B**). Because aging is relevant to pneumonia and the human cohort studied here, we also included aged mice (80-90 weeks old) challenged with saline, *Sp3*, or PR8. While all pneumonias were lethal at provided doses, the mice succumbed to bacterial infections more rapidly than viral infections (**Fig. 5C-D**). No differences in survival were observed between young and aged mice, likely because doses lethal even to young mice were used (**Fig. 5D**). All mouse samples were scored for the 20 histopathology features assessed in human samples. Some features were never observed in these acute pneumonia mouse models (abscess, desquamated epithelia, arterial thrombi, granuloma, hemosiderosis, hemophagocytosis, and peribronchiolar metaplasia). We were unable to assess microvessel fibrin due to technical limitations with the anti-fibrinogen antibody for mice. These missing features were excluded from further analyses.

**Figure 5.**
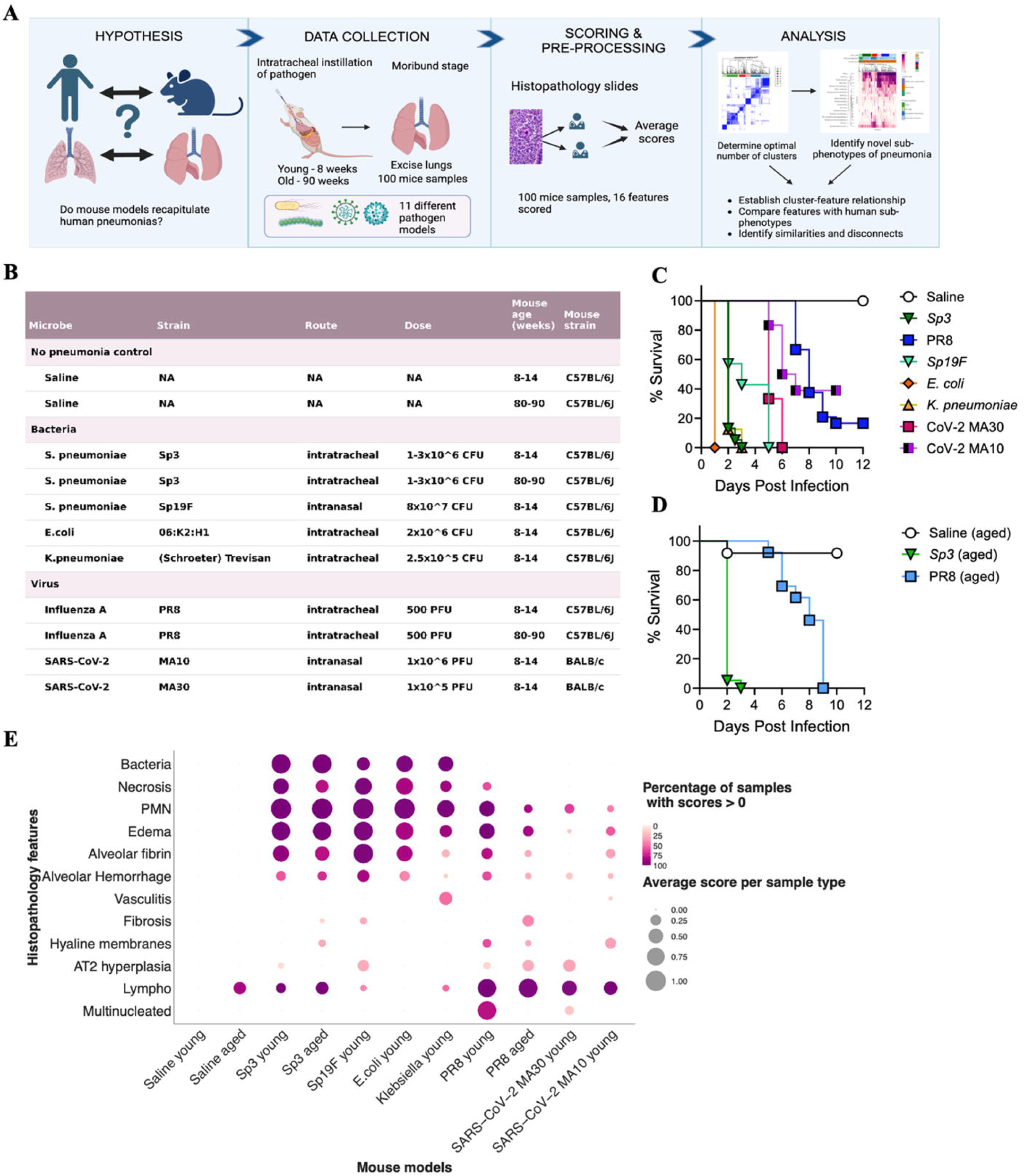
Heterogeneous pulmonary histopathologies in mouse models of pneumonia. (A) To examine patterns of pulmonary histopathology in mice with pneumonia, we utilized necropsy lung sections that were stained by H&E and anti-fibrinogen IHC before being scored by two pathologists, then analyzed computationally in the same manner as performed with the human pneumonia samples. (B) 11 mouse models using saline, bacteria, or viruses were used to obtain 100 murine pneumonia samples. Microbe strain, route and dose of infection as well as and mouse strain and age were consistent within each model. (C-D) Survival for each mouse was recorded across all models in young mice (in C) and for saline, Sp3, and PR8 in aged mice (in D). (E) Bubble plot of the variation in histopathology scoring for the 12 features scored in mice for each mouse model. Size of the bubble indicates average score per group and color indicates the percentage of samples within group with a non-zero score.

All of the mouse models of pneumonia elicited pulmonary histopathology that was absent from young mice receiving saline (**Fig. 5E, Supp. Fig. 7A**). Other than the presence of bacteria in sections, no histopathological feature completely distinguished viral and bacterial pneumonias (**Fig. 5E**). Neutrophils, necrosis, edema, and alveolar fibrin were prevalent and intense across bacterial infections, but more variable in viral infections (**Fig. 5E**). In contrast, lymphoplasmacytosis, multinucleated cells, and AT2 hyperplasia tended to be more prevalent and intense across viral infections (**Fig. 5E**). Fibrosis was most evident among aged mice with PR8 pneumonia (**Fig. 5E**) but not significant (**Supp. Fig. 7B**). Multinucleated cells were most prominent in young mice with PR8 pneumonia (**Fig. 5E, Supp. Fig. 7A**). Vasculitis was uniquely observed during *Klebsiella* pneumonia (**Fig. 5E, Supp. Fig 7A**).

### Mouse models of pneumonia involve heterogeneous pulmonary histopathologies

When the unsupervised machine learning framework previously applied to our human cohort was applied to our mouse cohort, 6 distinct clusters were identified (**Fig. 6A-B**; **Supp. Fig. 8**). Though clusters loosely grouped based on saline, viral, or bacterial pathogen, no samples from a single microbe were restricted to any one cluster (**Fig. 6A; Supp. Fig. 9**).

**Figure 6.**
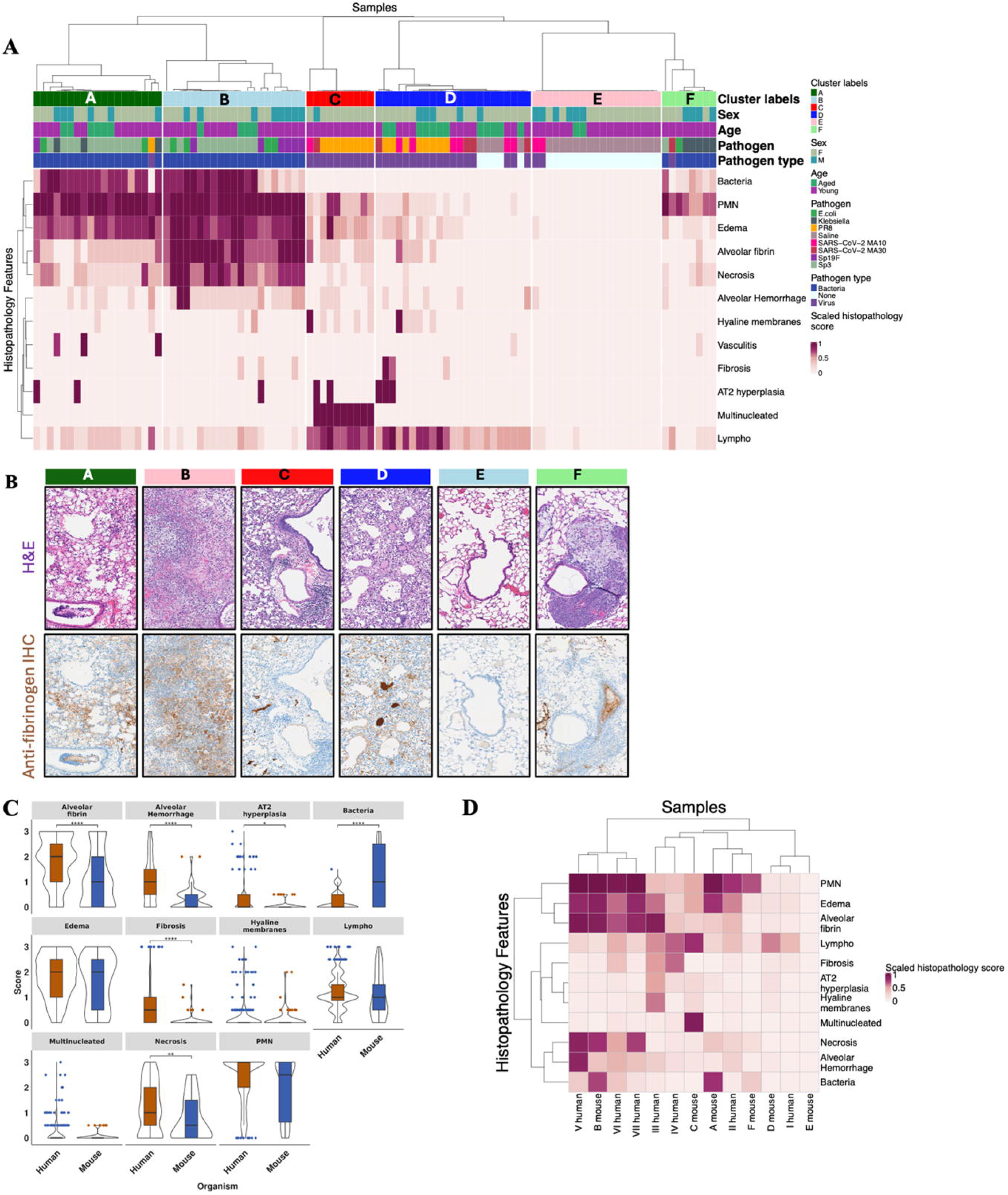
Mouse models reflecting human pneumonia histopathology sub-phenotypes. (A) Based on 12 scored features (rows) for each of 100 murine subjects (columns), clustering metrics identified 6 subsets of pneumonia (A-F) that were distinguished by pulmonary histopathology. (B) Representative images of H&E-stained and anti-fibrinogen IHC-stained sections for each cluster. (C) Histopathology features from human pneumonia samples were compared to mouse pneumonia samples across histopathology features detected in both species. Data were represented as median ± SEM and analyzed by Kruskal-Wallis test, followed by a post-hoc Dunn test and adjusted for multiple tests using Bonferroni correction method. (D) Mouse clusters segregated with human clusters with similar pulmonary histopathology. Aggregate profiles were generated by averaging feature scores per cluster and hierarchically clustering the combined dataset to show relationships between mouse clusters and human clusters.

Clusters A and B were characterized by high levels of bacteria, neutrophils, and edema, and cluster B additionally had abundant alveolar fibrin and necrosis (**Fig. 6A-B**). Nearly all mice in clusters A and B were infected by bacterial pathogens, without patterns specific to types of bacteria (**Fig. 6C, Supp. Fig. 9**). Samples in cluster C tended to have especially high lymphoplasmacytosis as well as multinucleated cells which were absent from other clusters (**Fig. 6A-B**); all of these samples were viral pneumonias in young mice, with PR8 representing 8 of the 10 samples (**Supp. Fig. 9**). Samples in cluster D were grouped primarily by high lymphoplasmacytosis with few other features present; these included young mice infected with SARS-CoV-2, aged mice infected with PR8, and some of the uninfected aged mice (**Fig. 6A-B, Supp. Fig. 9**). Samples in cluster E were grouped by the absence of pneumonia pathology and included mostly samples from saline-instilled mice (**Fig. 6A-B, Supp. Fig. 9**). Cluster F was primarily grouped by high neutrophilia with few other histopathology features; this contained most of the *K. pneumoniae*-infected lung samples (**Fig. 6A-B, Supp. Fig. 9**). As observed in humans, mouse models suggest distinct sub-phenotypes of pneumonia based on pulmonary histopathology. Mirroring the two large groups of human pneumonia presentations we observed in rapid autopsy analyses (**Fig. 2C**), the mouse bacterial pneumonias tended to have more neutrophils, edema, necrosis, alveolar fibrin, and alveolar hemorrhage, while mouse viral pneumonias tended to have more prominent lymphoplasmacytosis, fibrosis, and multinucleated cells.

### Connections between mouse models and human pneumonias

It is unclear how well mouse models of pneumonia match or disconnect with human pneumonia^35–39^. As noted above, 7 features observed in some lungs from human pneumonia patients were never observed in these mouse models. We compared the 11 histopathology features that were present in both our human and mouse pneumonia samples (omitting human “control” and mouse saline samples). The human pneumonia samples had significantly higher alveolar fibrin, alveolar hemorrhage, AT2 hyperplasia, fibrosis, and necrosis than the mouse models, while the bacterial mouse models had significantly higher amounts of bacteria observed in H&E-stained lung sections (**Fig. 6C**). Overall, these data suggest more severe and complex histopathology in human pneumonias, but more bacterial abundance in mouse models.

To determine relationships between the histopathology clusters observed in humans and in mice, we combined the 7 human and 6 mouse clusters into one dataset for integrated analyses. Human clusters V, VI, and VII and mouse cluster B segregated together, all of which had higher levels of neutrophils, edema, necrosis, and alveolar fibrin (**Fig. 6D**). Mouse cluster B was composed of samples from pneumococcus (*Sp3* and *Sp19F*) and *E. coli* (**Fig. 6A, Supp. Fig. 9**), suggesting that these bacteria in mice may yield necrosuppurative and hemorrhagic pneumonia sub-phenotypes. Human clusters III and IV segregated with mouse cluster C (**Fig. 6D**), including stronger signals for lymphoplasmacytosis, AT2 hyperplasia, and hyaline membranes. Mouse cluster C was primarily composed of PR8-infected samples (**Fig. 6A, Supp. Fig. 9**), suggesting that this acute mouse-adapted IAV infection captures some pneumonia features observed in these human pneumonia sub-phenotypes. Mouse clusters A and F segregated with human cluster II (**Fig. 6D**), characterized primarily by neutrophils and edema.

Mouse clusters A and F are composed of mostly bacterial pathogens, with *K. pneumoniae*-infected lung samples forming the majority of cluster F (**Fig. 6A, Supp. Fig. 9**); these infections in mice may model suppurative pneumonia. Finally, mouse clusters D and E segregated with human cluster I (**Fig. 6D**). These clusters included all the uninfected samples from both species (**Figs. 3A and 6A, Supp. Fig. 9**) plus some pneumonic human lungs in the “normal adjacent” sub-phenotype (**Fig. 3A**) and some viral pneumonias in both mouse clusters (**Fig. 6A, Supp. Fig. 9**). Altogether, these data reveal that mouse models capture much but not all of the pulmonary histopathology features and sub-phenotypes of pneumonia observed in human lungs.

## Discussion

This extensive analysis revealed and provided structure to diverse presentations of pulmonary pathology during pneumonia. Comparing pneumonic lungs to those with no known lung disease, we observed that some features anticipated to associate with pneumonia reached statistical significance, while others did not. This dataset also revealed that no single histopathological feature is required for a pathological diagnosis of pneumonia, since every feature was absent from some cases. Unsupervised clustering indicated that human pneumonias include at least 7 different sub-phenotypes, based on pulmonary histopathology.

Discrete sub-phenotypes also emerged from experiments using mice, and there was substantial but incomplete concordance between the naturally occurring pneumonias in humans and the experimental pneumonias in mice.

Considering the data from the human lung samples and the mouse models of pneumonia together, generalizations may emerge about etiology and histopathology. Across the human samples, features associated with each other to form two broad presentations, one which included more prominent neutrophils, edema, necrosis, alveolar fibrin, and alveolar hemorrhage, while the other included more prominent lymphoplasmacytosis, fibrosis, AT2 hyperplasia, peribronchiolar metaplasia, and multinucleated cells. The former associated with bacteria and the latter with SARS-CoV-2. In mouse models, the multiple bacterial infections resulted in greater neutrophils, edema, necrosis, and alveolar fibrin, while the multiple viral infections led to more prominent lymphoplasmacytosis, AT2 hyperplasia, and multinucleated cells. We propose that bacteria tend to result in suppurative, necrosuppurative, or hemorrhagic pneumonia sub-phenotypes, while viruses more often yield normal adjacent, DAD, or chronic interstitial pneumonia sub-phenotypes. These conclusions from systematic and unbiased analyses bolster current concepts of bacterial and viral pneumonia pathologies^40,41^.

One pneumonia sub-phenotype (cluster VI) was named “mixed” by us since it did not seem to reflect a single unified histopathology presentation. It is possible that this might represent multiple sub-phenotypes which the current analyses were not sufficiently empowered to fully differentiate. In addition, this cluster may include early (developing) and/or late (recovering) stages of other clusters. Furthermore, if etiological inferences are valid and viruses tend to yield clusters I, III, and IV while bacteria tend to yield clusters II, V, and VII, then perhaps cluster VI may include more complex etiologies such as secondary bacterial pneumonia after viral infection, combining different types of pulmonary pathologies.

When samples from human pneumonias and mouse pneumonias were considered together, the distinct histopathological presentations defined these infected lungs more than the different experimental settings or host species. These data support translational relevance and suggest that mouse models of pneumonia may be useful for dissecting mechanisms underlying pulmonary pathology sub-phenotypes observed in humans. These models thus may be used to elucidate the impact of host-directed therapies on outcomes during distinct pneumonia sub-phenotypes.

However, the match between human pneumonias and mouse models was imperfect. Of the 7 histopathology features observed in human lungs but never in mouse models, some were rare even in humans (hemophagocytosis, abscess, granuloma, and arterial thrombi) while others were found in a fifth to a half of human samples (peribronchiolar metaplasia, hemosiderosis, and desquamated epithelia). Such features may be later stages of pneumonia processes which are not captured in the acute mouse models examined here. This timing issue also may be a reason that other features were higher in humans compared to mice, such as fibrosis. Conversely, bacteria were more prevalent in mouse models than human pneumonias, which may result from antibiotics being often used in humans but never in these mouse studies. Hyaline membranes and AT2 hyperplasia were observed in mouse samples, but sparsely and never together; thus, the DAD sub-phenotype was not effectively captured with these model pneumonias. Relevant to these last two discrepancies, animal studies that include clinically relevant therapies (like antibiotics) and supportive measures (like mechanical ventilation) might increase concordance of sub-phenotypes across species^17^. Finally, mice in hyper-clean pathogen-free facilities are ineffective at capturing aspects of human immunology, which can limit applicability^42–44^. Introducing relevant microbial histories to mice seeds their lungs with resident memory T and B cells and trained alveolar macrophages, all of which make mouse lungs more “human-like”^45^. The mouse models used here capture many relevant aspects of human pneumonia, but ensuing studies should tailor experimental systems to address some of the limitations observed and make animal studies even more translational.

The biology underlying these sub-phenotypes needs now to be characterized, including kinetics, underlying mechanisms, and functional significance of the observed histopathologic features and sub-phenotypes. Future studies are indicated to determine whether host-directed therapies like corticosteroids, anticoagulants/fibrinolytics, biologics, and many others may have disparate effects for distinct sub-phenotypes of pneumonia. Since mouse models were useful in capturing pneumonia sub-phenotypes, they may empower next steps. Before these anticipated advances in biological knowledge can possibly inform clinical applications, it will also be necessary to determine if these sub-phenotypes can be ascertained using less invasive approaches than pulmonary histopathology, including blood sampling, bronchoscopy, radiology, exhaled breath analyses, or combinations of approaches.

The degree to which the pulmonary histopathology sub-phenotypes observed here are broadly relevant will need further exploration. Clustering algorithms are always influenced by the number of samples included, and increasing sample size could reveal additional sub-phenotypes. Our samples were collected from an elderly and predominantly white population from one location (greater Phoenix, AZ); the inclusion of younger patients from more diverse backgrounds and locales could reveal different pneumonia presentations. Autopsies are but a snapshot in time. Sub-phenotypes observed here may reflect different stages of biological processes, as with the hyperinflammatory and hypoinflammatory sub-phenotypes of ARDS^46^. For example, suppurative pneumonia may precede necrosuppurative and/or hemorrhagic pneumonias. Using autopsy samples may skew the dataset to over-represent more severe and life-threatening forms of pneumonia. The distribution of pneumonia cases amongst these 7 sub-phenotypes derived from an autopsy dataset should not be extrapolated to all settings of pneumonia, such as the more common situation when pneumonia is survived. Despite these inherent limitations, the concordance of human clusters with mouse clusters suggests that the pulmonary histopathology sub-phenotypes derived from these human subjects are observed across species, supporting generalizability.

Our study reveals that pneumonia histopathologies are diverse and heterogeneous, but cluster to form distinct sub-phenotypes in human samples and in mouse models. These results encourage further exploration of pneumonia sub-phenotypes, using additional sampling strategies (beyond autopsy lung samples) and analytic measures (beyond histopathology). The pneumonia sub-phenotypes that were derived from pulmonary histopathology suggest new directions of investigations to determine whether these different sub-phenotypes can be discriminated in living subjects, are driven by divergent biological processes, and respond differently to host-directed therapies.

## Materials and Methods

### Human rapid autopsy lung biobank

We utilized postmortem human lung tissues from retirement-age subjects in the greater Phoenix, AZ retirement communities who consented for research-purpose rapid autopsies between the years 2006-2020^29^ from the Brain and Body Donation Program of Banner Sun Health Research Institute. Approximately 13% of the enrolled subjects die each year with a median age of 82 years at autopsy. The median time from death until tissues are collected and processed is 3 hours, allowing for exceptional cell and molecular integrity. A total of 404 samples were analyzed and controlled for quality; duplicate samples from the same patients were removed. This retrospective source provided lung samples with complete autopsy records for 292 subjects: 276 samples were from patients given a diagnosis of pneumonia upon autopsy, 25 of whom were from patients with a COVID-19 diagnosis including a SARS-CoV-2 RT-PCR positive nasopharyngeal swab collected in 2020, plus another 16 samples from patients with no known lung disease based on autopsy and medical record.

### Human metadata

De-identified patient data was used to extract the metadata information. Age at the time of death was calculated using sample ID which consists of the year of death, and with self-reported date of birth. Self-reported smoking information was processed by grouping patients with reported terms (“smoke”, “smoking”, “tobacco”). Terms such as “marijuana” and “etoh” were not included as only reported rarely. Patients with missing entries or conflicting entries were marked as “not recorded.” Any patient with an entry of “none” for smoking was recoded as non-smoker. To classify former and current smokers, we used information about year in which a patient quit smoking. Patients who quit smoking 20 years before death were classified as “former smokers” and patient who quit smoking more recently (<20 years) before death were classified as “current smokers.” Patients with missing information or conflicting information were classified as “not recorded.” Similarly, alcohol status and history were determined using the self-reported alcohol terms. and patients with conflicting or missing terms were classified as “not recorded.” Other metadata variables such as binarized sex and race were generated using the self-reported information.

### Mice

C57BL/6J (#000664) and BALB/cJ (#000651) mice between the ages of 8-14 weeks and 80-90 weeks of age were obtained from the Jackson Laboratories (Bar Harbor, ME). Both male and female mice were included for all studies. Mice were housed in specific pathogen-free conditions on a 12-hour light-dark cycle with 30-70% humidity and 20-26°C temperature with access to standard mouse chow and water *ad libitum*. Mice were euthanized by overdose of anesthesia, inhaled isoflurane and exsanguination before organ collection in most cases, or ketamine/xylazine for the mice infected with MA10 and MA30 in biosafety level 3 laboratory (BSL-3) facilities. All SARS-CoV-2 mouse experiments were performed with BSL-3 precautions at the Boston University National Emerging Infectious Diseases Laboratories (BU-NEIDL), whereas all other mouse experiments were performed in BSL-2 laboratory space. Mouse experiments were performed following recommendations stated in the Guide for the Care and Use of Laboratory Animals of the National Institutes of Health^47^. The protocols were reviewed and approved by the Institutional Animal Care and Use Committee at the Boston University Chobanian and Avedisian School of Medicine.

### Pathogens and mouse infections

*Streptococcus pneumoniae* serotype 3 [ATCC 6303], *Escherichia coli* 06:K2:H1 [ATCC 19138], and *Klebsiella pneumoniae* serotype O1:K2 [ATCC 43816] were cultured on sheep’s blood-agar plates at 37°C for 10-12 hours. *S. pneumoniae* serotype 19F [EF3030] was cultured on sheep’s blood-agar plates at 37°C for 12-14 hours, then re-cultured onto sheep’s blood-agar plates at 37°C for 4-6 hours before dilution. Influenza A virus A/PR/8/34 (PR8), acquired from Dr. Nicholas Heaton (Duke University)^48^, was propagated in chicken eggs and infectious virus was measured by plaque assay on MDCK cells. Mouse-adapted coronavirus SARS-CoV-2 MA10 (NR-55329) was obtained from BEI resources, and MA30 was from Dr. Stanley Perlman (University of Iowa)^49^. Both MA10 and MA30 were propagated in VeroE6 cells and viral titer was measured by plaque assay in VeroE6 cells. Mice were anesthetized by intraperitoneal (i.p.) injection of a ketamine/xylazine injection (or 1-3% isoflurane for SARS-CoV-2 infections) and infected either intranasally (i.n.) or intratracheally (i.t.) by surgically exposing the trachea and instilling 50 µl of each pathogen diluted in sterile saline or 1XPBS via a 24-gauge angiocatheter directly into the left lung lobe. Mouse weights and clinical disease were monitored daily. For all mice infected with pathogens other than SARS-CoV-2, mice were euthanized when they fulfilled 1 or more moribund criteria, including weight loss greater than 20% (bacterial infections) or 30% (viral infections) of their initial weight, labored breathing, altered gait, and/or extreme lethargy.

For SARS-CoV-2 MA10 and MA30 experiments, a clinical scoring system defined disease progression. The score of “1” was given for each of the following situations: body weight, 10– 29% loss; respiration, rapid and shallow with increased effort; appearance, ruffled fur and/or hunched posture; responsiveness, low to moderate unresponsiveness; and neurological signs, tremors. The sum of these individual scores constituted the final clinical score (0-5). Mice were humanely euthanized if they were moribund by the above characteristics or if they received a clinical score of 4 or above for two consecutive days.

### Histopathology scoring of human and mouse lung samples

Formalin-fixed, paraffin-embedded (FFPE) human lung sample blocks were sectioned to 5μm and stained by H&E and IHC using a rabbit anti-fibrinogen gamma chain antibody (CST 45850S, #E1U3Z). FFPE mouse lung samples were similarly sectioned and stained by H&E and IHC using a rabbit anti-fibrinogen gamma chain antibody (Abcam ab281924, #EP24539-127). All staining was performed using a ST5010 linear autostainer (Leica). Histopathology was independently scored for each sample by two board-certified pathologists (NAC and DGR) for 20 distinct features (**Supp. Fig. 1A**) with a score range from 0-3, with 0 indicating a feature was not present in the section, 1 if the feature was present in <5% of the section, 2 if the feature was present in 5-25% of the section, and 3 if the feature was present in >25% of the section. Scores from the two pathologists were compared, and differences and means were calculated. If scores differed by 2 or more, slides were re-analyzed before calculating mean.

### Quantitative multiplex fluorescence IHC (qmf-IHC)

Out of 292 human lung samples, a subset of 159 were of sufficient quality to quantify leukocyte signals using qmf-IHC. Samples were stained using a Ventana Discovery Ultra (Roche) tissue autostainer using the following antibodies from Cell Signaling Technology and TSA-labeled fluorescent dyes from Akoya Biosciences, listed in sequential order of staining: MPO (#14569, clone E1E7I) and Opal 570 (FP1488001KT), CD19 (#90176, clone D4V4B) and Opal 480 (FP1500001KT), CD8a (#85336, clone D8A8Y) and Opal 620 (FP1495001KT), CD68 (#76437, clone D4B9C) and Opal 690 (FP1497001KT), and CD4 (#48274, clone EP204) and Opal 520 (FP1487001KT), with optimized staining conditions for this panel detailed in **Supp. Table 1**. Antigen retrieval was conducted using a Tris based buffer-CC1 (Roche). Fluorescently-labeled slides were imaged using a PhenoImager^TM^ Quantitative Pathology Imaging System (Akoya Biosciences, Marlborough, MA). Exposures for all Opal dyes were established on regions of interest with strong signal intensities to minimize exposure times and maximize the specificity of the signal detected. To maximize signal-to-noise ratios, images were spectrally unmixed using a synthetic library specific to each Opal fluorophore and DAPI. Furthermore, an unstained section was used to create an autofluorescence signature that was subsequently removed from whole-slide images using InForm software version 2.4.8 (Akoya Biosciences, Marlborough, MA). InForm exported the multispectrally unmixed images as QPtiffs, which were then fused together as a single whole slide image in HALO (Indica Labs, Albuquerque, NM) for image analysis.

For quantifying the area of the slide that contained MPO and CD68, the HALO (Indica Labs, Albuquerque, New Mexico) Area Quantification (AQ) module (v2.1.11) was created and finetuned to quantify the immunoreactivity based on color and stain intensity. This algorithm outputted the fraction of total area displaying immunoreactivity across the annotated whole slide scan in micrometers squared (μm²). For quantifying the absolute number and overall percentage of cells containing CD4, CD8, or CD19 staining, we utilized the Halo (Indica Labs) HighPlex phenotyping modules (HP, v4.0.4). In brief, this algorithm was used to first segment all cells within the annotated lung sections using DAPI counterstain. Next, minimum cytoplasm and membrane thresholds were set for each fluorophore to detect positive staining within each of the segmented cells. Parameters were set using the real-time tuning mechanism that was tailored for each individual sample based on signal intensity. The algorithm yielded outputs of total cell counts of CD4+, CD8+, and CD19+ cells and the total area analyzed. The quantitative output for AQ and HP were exported as a .csv file for downstream statistical analysis.

### Clustering of histopathology features

Histopathology feature scores were standardized across samples using min-max scaling (i.e., subtracting the minimum and dividing by the maximum) to obtain values between 0 and 1. We employed consensus clustering^50^ with the partition amongst medoids (PAM)^51^ algorithm using Euclidean distance to cluster the samples. All features were considered for each subset iteration with sub-sampling set of samples to 80%. Along with the average item consensus score for each cluster at each partition, we used other variance and information-based metrics such as the Bayesian Information Criterion (BIC) from the mclust function^52^, dunn2 index, dunn index, ch index, entropy and average silhouette width^53^ to determine the optimal number of clusters. The same process for clustering and determining the optimum number of clusters was used to analyze both human and mouse samples.

### Variable importance and decision tree analysis

Variable importance is often used in supervised learning to explain the features responsible for classification. However, clustering lacks ground truth labels and relies upon methods such as differential expression tests (Wilcoxon or ward test) to determine features different between clusters. We assumed the clusters generated by consensus clustering as the ground truth and used the random forest algorithm from R package randomForest to determine the variable importance for each cluster. All samples (min-max scaled) were used with random forest since the goal was not to train the model but to understand the path of clustering.

### Statistical tests

All pairwise group-level comparisons for histopathology features were performed using the Kruskal-Wallis test followed by the posthoc Dunn test (‘rstatix’ package). Scores were adjusted for multiple hypothesis testing using the Bonferroni method. All statistical test results are reported under supplementary data files (**Supp. Data file 1, Supp. Data file 2, Supp. Data file 3**). All data analyses were performed using R programming language, with scripts available on GitHub (https://github.com/campbio-manuscripts/Pneumonia_Subtypes). Correlations were generated using Spearman correlations. All pairwise comparisons for immunohistochemistry features were performed using Wilcoxon test and adjusted for multiple hypothesis testing using the Bonferroni method. Enrichments are computed using Fishers exact test.

### Other tools and Packages

All analysis was done using R programming language version 4.2.1. Visualization was generated using complexheatmap and ggplot2 packages. All cleaning and processing were done using the dplyr and tidyverse packages. More detailed information on packages used were provided as session information in Supp. Data table 4.

## Acknowledgments

This research was supported by grants from the NIH (R01 AI162850 to J.P.M., R01 HL171499 to J.P.M., R01 AI115053 to J.P.M., T32 HL007035 to J.P.M., F32 HL170650 to B.E.H., and R01 HL158732 to K.E.T.). Processing of histology tissues, H&E staining, anti-fibrinogen IHC staining, qmf-IHC staining and analysis, and microscopy imaging was performed through the BU-NEIDL Comparative Pathology Laboratory (NCPL) utilizing NIH funded S10 instrumentation (S10 OD030269 & S10 OD026983 to N.A.C). We are grateful to the Arizona Study of Aging and Neurodegenerative Disorders/Brain and Body Donation Program at Banner Sun Health Research Institute, Sun City, Arizona, for the provision of human materials, which has been supported by the NINDS (U24 NS072026 National Brain and Tissue Resource for Parkinson’s Disease and Related Disorders), NIA (P30 AG019610 and P30 AG072980, Arizona Alzheimer’s Disease Center), the Arizona Department of Health Services (contract 211002, Arizona Alzheimer’s Research Center), the Arizona Biomedical Research Commission (contracts 4001, 0011, 05-901 and 1001 to the Arizona Parkinson’s Disease Consortium), and the Michael J. Fox Foundation for Parkinson’s Research.

---

**Supplemental Figure 1. Histopathology features defined in human rapid autopsy samples.** (A) 20 pneumonia histopathology features were scored in human rapid autopsy samples, including the abbreviation used in presented figures and indication whether each feature was included in the analyses presented. (B-L) Representative images of the 11 least common pneumonia histopathology features observed in human samples, including abscess (B, H&E), arterial thrombi (C, H&E and anti-fibrinogen IHC), AT2 hyperplasia (D, H&E), bacteria (E, H&E), granuloma (F, H&E), hemosiderosis (G, H&E), hyaline membranes (H, H&E and anti-fibrinogen IHC), multinucleated cells (I, H&E), peribronchiolar metaplasia (J, H&E), vasculitis (K, H&E), and hemophagocytosis (L, H&E).

**Supplemental Figure 2. Scored features that did not differ depending on SARS-CoV-2 or visible bacteria status.** (A-B) Histopathology scores from pneumonia samples with a SARS-CoV-2 diagnosis (A) or with visible bacteria in the H&E section (B) were compared to all other pneumonia samples. Data are represented as median ± SEM and analyzed by Kruskal-Wallis test, followed by a post-hoc dunn test and adjusted for multiple tests using Bonferroni correction method.

**Supplemental Figure 3. Computational metrics guiding optimum cluster numbers in human dataset.** (A) Various variation-based and distance-based metrics such as average silhouette width, Calinski and Harabasz (ch) index, within cluster sum of squared distances (k-means objective function), dunn index, dunn2 index, ratio between within cluster sum of squared and between cluster sum of squared distances (wb.ratio), separation index (sindex). (B) Bayesian Information Criterion (BIC) shows the optimum partition to be at seven, with high BIC score and VEV as the best gaussian mixture model our data. (C) Consensus score for each pair of samples were calculated using the ratio of number of times each pair ended up in the same cluster to number of samples subsampled together. The heatmap shows the consensus score (high score is colored by dark blue and low score is colored by white) for the dataset at K=6, K=7 and K=8. (D) By using cluster labels as ground truth, a random forest classifier was trained and variable importance (VI) scores were calculated. Variable importance score highlights the top 6 features per cluster driving the cluster separation. The color of the dots indicate whether the feature had a higher or lower average score compared to samples in other clusters.

**Supplemental Figure 4. Histopathology scores across each metadata variable, including smoking and alcohol status, age, and sex.** (A-D) Histopathology scores from pneumonia samples are compared across binarized age groups (A), sex (B), smoking status (C), and alcohol drinking status (D). Data are represented as median ± SEM and analyzed by Kruskal-Wallis test, followed by a post-hoc dunn test and adjusted for multiple tests using Bonferroni correction method.

**Supplemental Figure 5. Pneumonia sub-phenotype is not influenced by available metadata variables including smoking and alcohol status, age, and sex.** (**A-D**) Samples were binarized for Age (A) as group 1 (43-69 years) and group 2 (70 -103), sex (B), smoking status (C) and alcohol drinking status (D) and cluster composition was examined across all 7 clusters.

**Supplemental Figure 6. Immunostaining comparisons across different conditions.** Immunofluorescence intensities were compared between control samples and pneumonia (A), binarized SARS-CoV2 status (B) and binarized visible bacteria in the lung (C) Data are represented as median ± SEM and analyzed by two-tailed Wilcoxon test and adjusted for multiple tests using Bonferroni correction method.

**Supplemental Figure 7. Mouse models differ in histopathology features compared to saline.** We compared the histopathology features across mouse models by comparing it to hanging controls saline young (A) and saline aged (B). Data are represented as median ± SEM and analyzed by two-tailed Wilcoxon test and adjusted for multiple tests using Bonferroni correction method.

**Supplemental Figure 8. Computational metrics guiding optimum number of clusters in mouse dataset.** (A) Various variation-based and distance-based metrics such as average silhouette width, Calinski and Harabasz (ch) index, within cluster sum of squared distances (k-means objective function), dunn index, dunn2 index, ratio between within cluster sum of squared and between cluster sum of squared distances (wb.ratio), separation index (sindex). (B) Bayesian Information Criterion (BIC) shows the optimum partition to be at six, with high BIC score and EEV as the best gaussian mixture model our data. (C) Consensus score per each pair of samples were calculated using the ratio of number of times each pair ended up in the same cluster to number of samples subsampled together. The heatmap shows the consensus score (high score is dark blue and low is white) for the dataset at K=5, K=6 and K=7.

**Supplemental Figure 9. Proportions of different mouse models across clusters show heterogenous cluster membership.** Samples were grouped based on the pathogen used to infect the mice and proportions were calculated for each cluster.

**Supplemental Table 1. Optimized conditions for qmf-IHC panel staining.**

